# Emergence of multifrequency activity in a laminar neural mass model

**DOI:** 10.1101/2025.06.16.659878

**Authors:** Raul de Palma Aristides, Pau Clusella, Roser Sanchez-Todo, Giulio Ruffini, Jordi Garcia-Ojalvo

## Abstract

Neural mass models (NMMs) aim to capture the principles underlying mesoscopic neural activity representing the average behavior of large neural populations in the brain. Recently, a biophysically grounded laminar NMM (LaNMM) has been proposed, capable of generating coupled slow and fast oscillations resulting from interactions between different cortical layers. This concurrent oscillatory activity provides a mechanistic framework for studying information processing mechanisms and various disease-related oscillatory dysfunctions. We show that this model can exhibit periodic, quasiperiodic, and chaotic oscillations. Additionally we demonstrate, through bifurcation analysis and numerical simulations, the emergence of rhythmic activity and various frequency couplings in the model, including delta-gamma, theta-gamma, and alpha-gamma couplings. We also examine how alterations linked with Alzheimer’s disease impair the model’s ability to display multifrequency activity. Furthermore, we show that the model remains robust when coupled to another neural mass. Together, our results offer a dynamical systems perspective of the laminar NMM model, thereby providing a foundation for future modeling studies and investigations into cognitive processes that depend on cross-frequency coupling.

**Author Summary:** Understanding how the brain generates and coordinates rhythms across different layers is essential for uncovering the mechanisms underlying perception, memory, and cognition. In this work, we analyze a previously developed model of mesoscopic brain activity that simulates the layered structure of the cortex and its ability to produce coupled slow and fast neural oscillations. Using tools from dynamical systems theory, we reveal how the model gives rise to a rich repertoire of dynamical patterns—including periodic, quasiperiodic, and chaotic activity—through the coexistence of multiple oscillatory modes. We also investigate how pathological changes, such as those linked to Alzheimer’s disease, alter the model’s dynamics and impair its capacity to sustain complex cross-frequency interactions. Finally, we show that the model remains stable when connected to another brain region, highlighting its robustness. Our findings provide a deeper understanding of how multifrequency neural rhythms may emerge, how they might break down in disease, and how this modeling framework can inform both future theoretical studies and the development of new brain models.

## 1 Introduction

Oscillations are ubiquitous in the brain [1]. These rhythmic or repetitive patterns of electrical activity emerge from the firing of individual neurons, reflecting the synchronized activity of neuronal ensembles, and are therefore the result of the interplay between different scales: microscopic (single neurons), mesoscopic (local groups of neurons), and macroscopic (across brain regions) [2, 3]. The frequencies in which mammalian neuronal networks oscillate span four orders of magnitude, ranging from 0.05 Hz to 500 Hz, with this diversity arising from neuron and synapse properties, their interactions, and the circuit motifs that they form [4].

Neural mass models (NMMs) can capture the mesoscopic dynamics of ensembles of neurons as nonlinear oscillators [5–9]. In this formalism, a population is typically characterized by its average firing rate, membrane potential, as well as the characteristics of synapses through which it connects to other populations (time constant and amplitude of post-synaptic potentials, or PSPs). Although limited, these models have been instrumental in advancing our understanding of brain rhythms, successfully replicating brain activity patterns observed through electrophysiological and neuroimaging techniques [10–23].

In this paper we focus on the emergence of multifrequency coupling in the recently introduced *laminar* NMM (LaNMM) [24], an extended NMM that includes the layered structure of the cortex and describes the frequency specificity and coupling between layers. Such a phenomenon appears to be crucial for brain activity, and plays a role in cognitive functions [25–32]. Recent work with the LaNMM has shown how this model can be used to implement gating and predictive coding elements [33] and cooperation/competition across brain regions [34]. Frequency specificity is observed across cortical layers, with different frequency bands contributing to distinct aspects of neural processing and communication [26, 35, 36]. For example, during a memory task, macaques exhibit high-frequency activity in superficial layers and low-frequency activity in deep layers. Moreover, the phase and amplitude of deep-layer low-frequency oscillations influence the phase and amplitude of superficial gamma dynamics, highlighting the intricate interplay between frequencies across cortical layers [36].

From the dynamical point of view, this is the result of the interaction of different nonlinear oscillators, each one with a different natural frequency. In turn, this frequency depends on the intrinsic properties of each neuronal population, such as the synaptic characteristics of neurons that make up the population and the connectivity between them. Hence, when we consider the interaction between different populations, we have to take into account the tendency of each population to oscillate in its preferred frequency. From this interplay, many interesting and complex phenomena emerge, including, multifrequency coexistence, chaos and cross-frequency coupling [37–42].

The LaNMM combines two well-known NMMs: the Jansen and Rit’s (JR) model [7, 8, 43] and a model implementing the pyramidal-interneuron network gamma (PING) mechanism [5, 44–47]. The first model, consisting of a population of pyramidal neurons coupled to two other populations, is responsible for sustaining lower frequencies and is associated with deeper cortical layers. The latter model includes a pyramidal and an interneuron population, and generates faster rhythms in the gamma band. Finally, inputs to the two pyramidal populations are added to simulate signals from other cortical regions. By integrating these elements, the LaNMM elegantly displays cortical oscillations across different frequency bands [22, 24], and has been used to model phenomena including effects of serotonergic psychedelics [48], oscillatory changes in Alzheimer’s disease [33, 49], predictive coding mechanisms [33], and cooperative vs. competitive interactions [34]. In particular, the last two studies build on the cross-frequency coupling characteristics of the LaNMM.

Here, we analyze the LaNMM’s ability to display multifrequency dynamics as a function of the (constant) inputs of its pyramidal populations. Specifically, we conduct a bifurcation analysis of the model with respect to changes in the input parameters. We show that in addition to the alpha-gamma coupling described in [22, 24, 33, 34], the model is also capable of generating delta-gamma and theta-gamma coupling, which, to our knowledge, has not been previously reported for other NMMs. Our analysis reveals that the parameter space of external inputs contains a large region of quasiperiodicity, where multifrequency coupling emerges due to the coexistence of two unstable limit cycles associated with the two components of the model: the JR and the PING. Additionally, we examine how alterations associated with Alzheimer’s disease impair this capability, by analyzing their impact on the bifurcation diagram and how they reshape the regions where the system exhibits multifrequency activity. Furthermore, we demonstrate that the model remains robust when coupled to other neural masses.

## 2 Results

### 2.1 Laminar Neural Mass Model

Neural mass models are lumped mathematical descriptions of neuronal activity that aim to mimic the behavior of populations of neurons with a small number of differential equations. Our main object of study is the LaNMM, first proposed in [22, 24], which consists of five interconnected populations of neurons, each representing a different cell type:

- Pyramidal cells located at deep cortex layers (*P*_1_).
- Pyramidal cells located at superficial layers (*P*_2_).
- Fast inhibitory parvalbumin-positive cells (*PV*).
- Slow inhibitory somatostatin-expressing cells (*SST*).
- Excitatory spiny stellate cells (*SS*).

Additionally, we consider two generic external inputs targeting *P*_1_ and *P*_2_ that represent incoming inputs from other neuronal populations. Figure 1 illustrates the model and the connectivity between these populations. Excitatory synapses, mediated by AMPA neurotransmitters, are represented as triangles, while inhibitory synapses, mediated by GABA neurotransmitters, are shown as circles.

**Figure 1.**
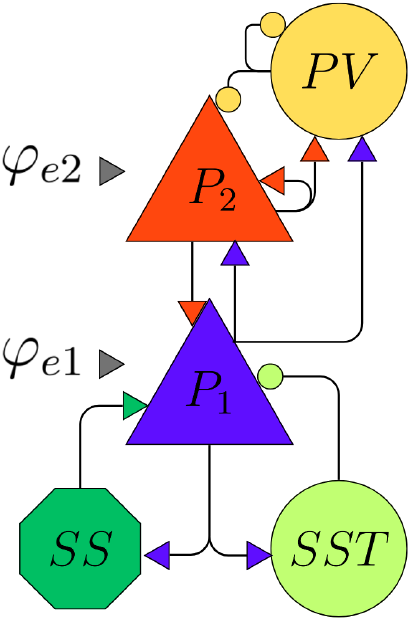
Illustration of the neuronal populations of the LaNMM and the connectivity between them. The top two blocks correspond to the PING model, and the bottom three blocks to Jansen and Rit’s model. Rounded shapes represent interneuron populations, while the rest represent excitatory populations. External inputs targeting *P*_1_ and *P*_2_ are represented by *φ*_*e*1_ and *φ*_*e*2_, respectively.

The dynamics of each population and their interactions are modeled following heuristic principles proposed by seminal studies in the field [5, 7, 50, 51]. First, the average axonal pulses originated in one population (with average firing rate *r*(*t*)) produce an average postsynaptic potential (PSP) *y*(*t*) via a linear convolution [51]:

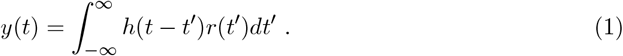

The kernel *h* reads:

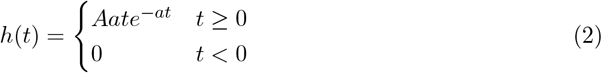

were parameters *A, a* vary depending on the effect of the modeled neurotransmitter on the postsynaptic population. For instance, while excitatory neurotransmitters cause excitatory postsynaptic potentials (EPSPs), which depolarize the postsynaptic population, inhibitory neurotransmitters cause inhibitory postsynaptic potentials (IPSPs), which have the opposite effect. Specifically, the amplitude and timescale of the postsynaptic potentials depend on the parameters *A*(mV), which represents the synaptic gain, and *a*(s^−1^), which is the time constant of the average PSP. Figure 2(a) shows the three different PSPs captured by Eq. (2) for individual populations (i.e. in the absence of coupling), one for excitatory synapses (mediated by AMPA neurotransmitter, *A*, blue line) and the other two for inhibitory synapses (mediated by slow and fast GABA neurotransmitter, *G*_*s*_ and *G*_*f*_, green and yellow lines respectively).

**Figure 2.**
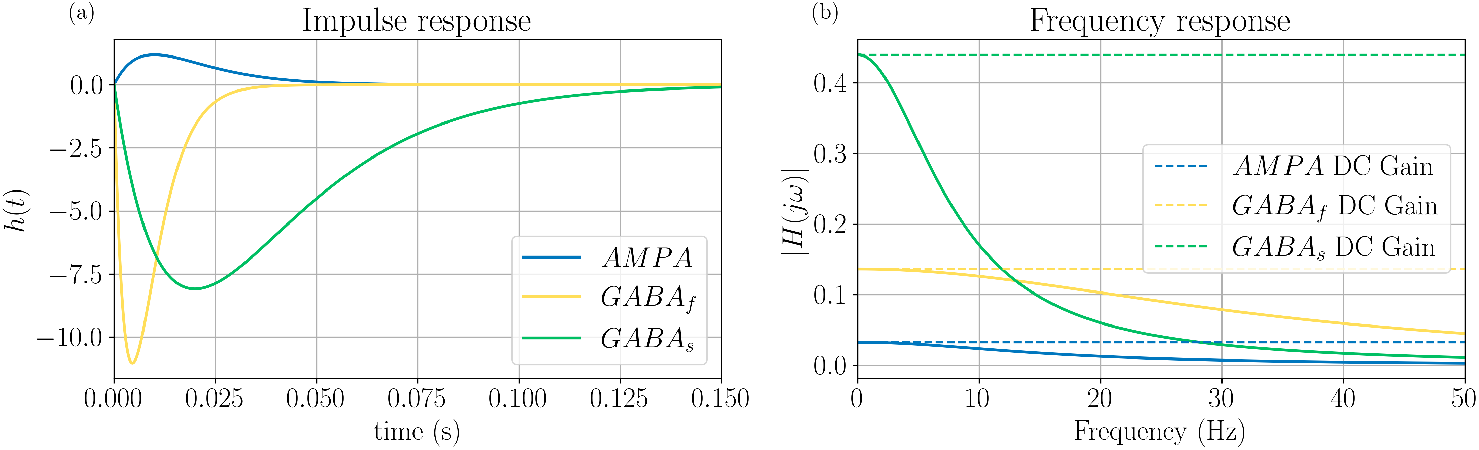
The three types of *h* kernels present in the LaNMM lead to different post-synaptic potentials. The PSP amplitudes and decay time are governed by the parameters *A* and *a*, respectively. The right hand side displays the frequency response of the filters.

In Fig. 2, the strongest PSPs are generated by GABAergic interneurons, with PV neurons (with *GABA*_*f*_) responding the fastest and SST neurons (with *GABA*_*s*_) the slowest. The IP-SPs have a larger amplitude than the EPSPs, due to inhibitory neurons establishing synapses closer to the cell body of postsynaptic cells [7, 52]. The frequency response associated with each h-block is shown in Fig. 2 (b), and is modeled using the expression 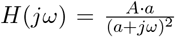, which describes how each synapse amplifies or attenuates signals across different frequencies. AMPA is the least responsive of all synapses, with a weaker response at higher frequencies. In contrast, fast GABA shows more responsiveness at higher frequencies, i.e. it is more sensitive to fast oscillations and rapid signals, which is typical for fast inhibitory synapses.

A second key ingredient of the modeling is a nonlinear function *σ*(*v*) that relates the average membrane potential (*v*) to an average firing rate [53]:

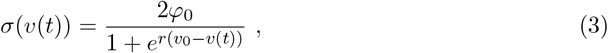

where *φ*_0_ is the half maximal firing rate of the targeted population, *v*_0_ is the value of the potential when *φ*_0_ is achieved, and *r* determines the steepness of the sigmoid at the threshold (*v*_0_, *φ*_0_).

Combining these concepts, the average membrane potential *y*(*t*) of a neuronal population (modulated by interactions with another population through the average membrane potential of the former *v*(*t*)) is governed by the following second-order differential equation [54] that implements the linear convolution described in Eq. 1:

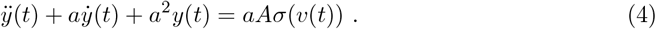

Using Eq.(4) and considering the connectivity scheme described in Fig. 1, the system of equations governing the model is given by

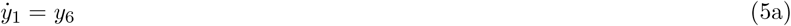

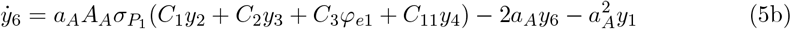

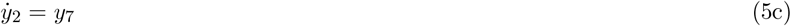

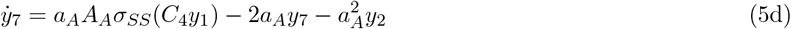

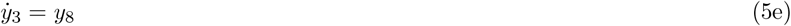

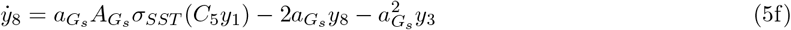

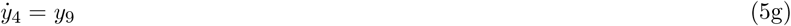

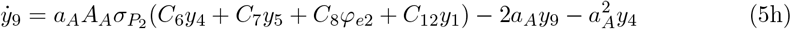

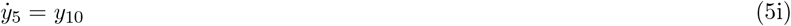

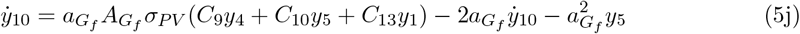

where the description and values of the parameters are given in Table 1. In this paper we will pay particular attention to the effects that the external inputs *φ*_*e*1_ and *φ*_*e*2_ (acting upon the pyramidal populations *P*_1_ and *P*_2_, respectively) have on the dynamics of the system.

**Table 1:** Description of model parameters and their standard values, as used in [22].

| Parameter | Description | Value |
| --- | --- | --- |
| $A_s$ | Average synaptic gain | $A_A = 3.25\text{mV}; A_{G_s} = -22\text{mV}$<br>$A_{G_f} = -30\text{mV}$ |
| $a_s$ | Time rate constant of average postsynaptic potentials | $a_A = 100\text{s}^{-1}; a_{G_s} = 50\text{s}^{-1}$<br>$a_{G_f} = 220\text{s}^{-1}$ |
| $C_s$ | Average number of synaptic contacts between population types | $C_1 = 108, C_2 = 33.7, C_3 = 1, C_4 = 135,$<br>$C_5 = 33.75, C_6 = 70, C_7 = 550, C_8 = 1, C_9 = 200,$<br>$C_{10} = 100, C_{11} = 80, C_{12} = 200, C_{13} = 30.$ |
| $v_0$ | Potential when 50% of the firing rate is achieved | 6mV (except for $P_2$: 1mV). |
| $\varphi_0$ | Half of the maximum firing rate | 2.5 Hz |
| $r$ | Slope of the sigmoid function at $v_0$ | $0.56\text{mV}^{-1}$ |
| $\varphi_e$ | External input | $\varphi_{e1} = \frac{A_3}{a_3} 200, \varphi_{e2} = \frac{A_8}{a_8} 90$ |

In the next section we present a detailed analysis of the dynamics and bifurcations of system (5).

### 2.2 Baseline dynamics of the LaNMM

In this section, we explore the dynamical properties of the system described by Eq. (5). We begin by analyzing the model’s response to external inputs. Specifically, we simulate the system starting from a fixed point (with *φ*_*e*1_ = *φ*_*e*2_ = 0) and apply a pulse simultaneously to both *P*_1_ and *P*_2_ at *t* = 0. Figure 3 illustrates the time evolution of the membrane potentials of the model’s two main neural populations. These observables are given by [22]:

**Figure 3.**
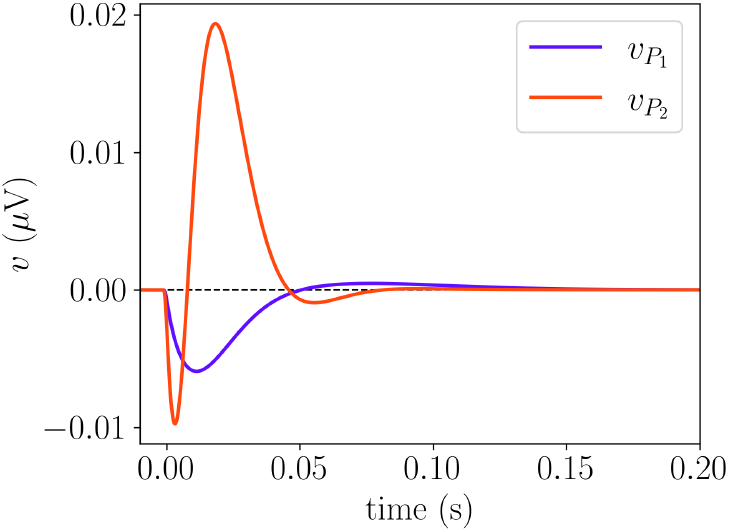
Response of the model for a pulse of 1 ms delivered to *P*_1_ and *P*_2_ simultaneously. Parameters values as in Table 1, with *φ*_*e*1_ = 0 and *φ*_*e*2_ = 0.

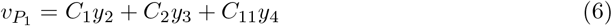

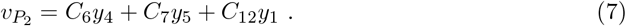

For better visualization, we center the signals at *v* = 0 by removing their DC components, to enhance the comparison between responses. We observe differences not only in amplitude but also in the timescale of the response. *P*_2_ exhibits a stronger response, characterized by a fast initial peak followed by damped oscillations back to the steady state. In contrast, *P*_1_ has a significantly slower response with a smaller amplitude. These differences in response dynamics align with the intrinsic properties of these populations: *P*_2_ has higher excitability (lower *v*_0_) and faster intrinsic time scales due to its coupling with *PV* cells. The slower recovery of *P*_1_ is influenced by *SST* cells, which exhibit the slowest response, as shown in Fig. 2. On the other hand, the network responses shown in Fig. 3 differ significantly from those observed in Fig. 2, which is to be expected since populations with distinct amplitudes and time constants now interact. We will revisit this response when analyzing the stability of the fixed point of the system in the next section.

As previously mentioned, a key feature of this neural mass model is its ability to sustain distinct oscillatory frequencies across different populations, particularly in the alpha and gamma bands. This behavior is illustrated in Fig. 4, where the time evolution of the membrane potentials of populations *P*_1_ and *P*_2_ are shown in panels (a) and (b), respectively, for *φ*_*e*1_ = 200 and *φ*_*e*2_ = 90 as in the model in [22]. While both populations exhibit rhythmic activity, their temporal dynamics differ significantly, with *P*_1_ oscillating at a lower frequency, around 10 Hz (alpha range), compared to the faster oscillations of *P*_2_, around 40 Hz (gamma range).

**Figure 4.**
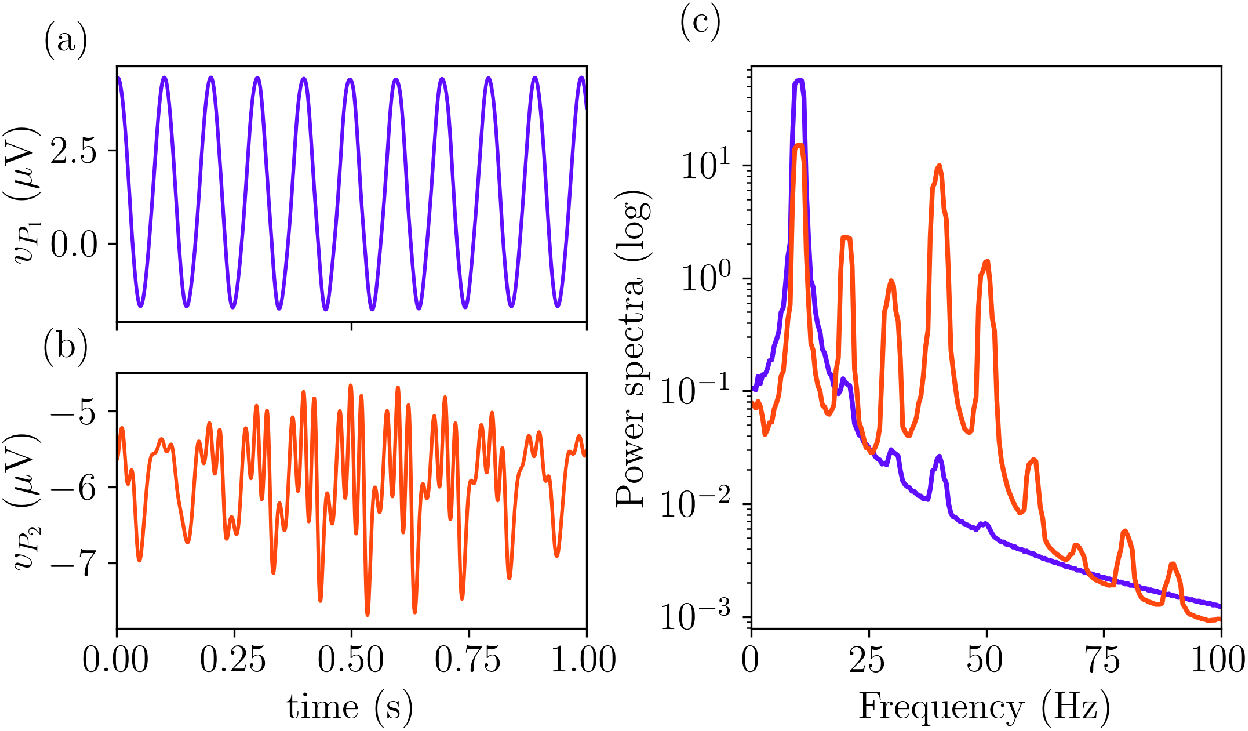
Time evolution of the membrane potentials of *P* 1 (a) and *P*_2_ (b) with the respective power spectra shown in panel (c). Parameters values as in Table 1 with *φ*_*e*1_ = 200 *φ*_*e*2_ = 90.

To further characterize these differences, panel (c) of Fig. 4 displays the power spectra of each signal. The two curves are plotted using the same colors as the corresponding time series in panels (a) and (b). This analysis confirms that while both populations share a strong *α* component around 10 Hz, *P*_2_ exhibits pronounced *γ*-band activity at around 40 Hz. The shared *α* rhythm enables cross-frequency coupling, where *γ* oscillations in *P*_2_ may be modulated by the phase of the *α* rhythm. These features were explicitly introduced into the model to ensure a representation of experimental results [22].

### 2.3 Fixed points and bifurcations

We now look for the fixed points of the system (5). To solve this system and find the fixed points, we used the continuation and analysis software AUTO-07p [55]. Figures 5(a) and (b) show a one-parameter bifurcation diagram obtained by varying the external input of the pyramidal population *P*_1_, *φ*_*e*1_, while keeping *φ*_*e*2_ = 0, in terms of the observables 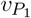 (a) and 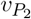 (b).

**Figure 5.**
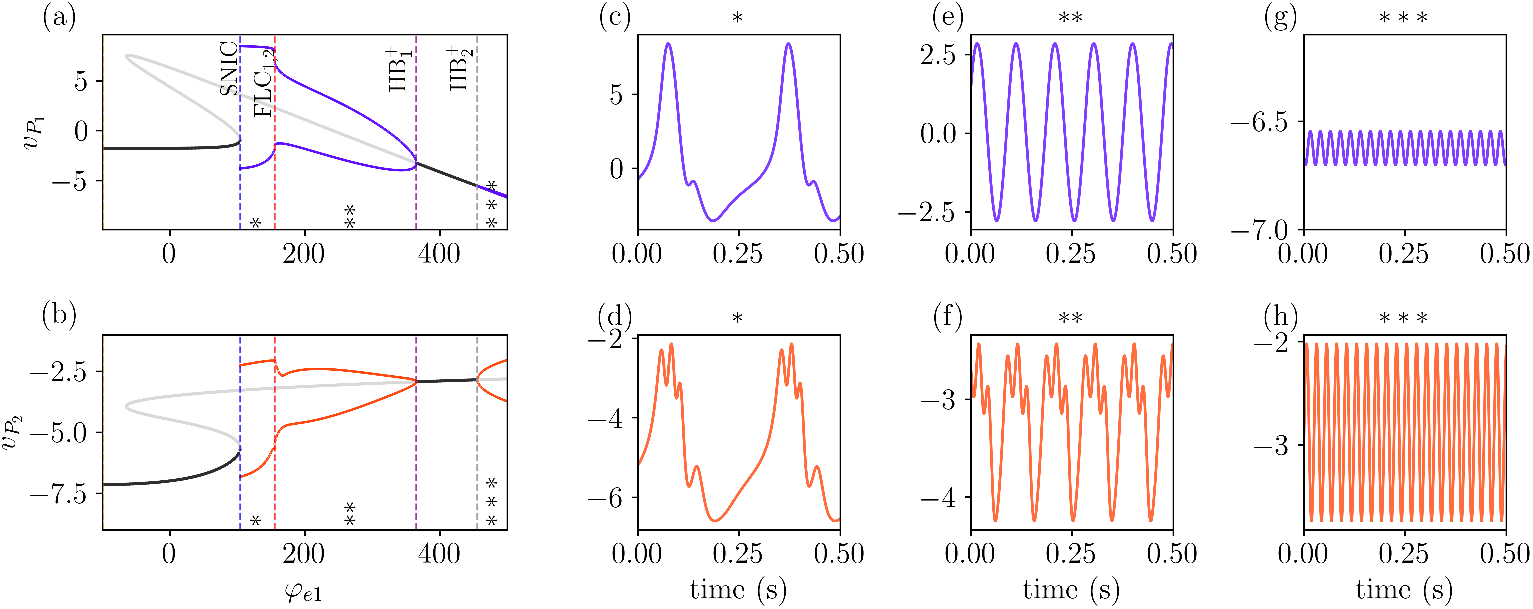
Bifurcation diagram of the system showing the fixed points and the amplitude of oscillatory solutions for *v*_*P* 1_ and *v*_*P* 2_ in panels (a) and (b), with *φ*_*e*2_ = 0. Stable (unstable) fixed points are shown in dark (light) grey. Increasing the external input (*φ*_*e*1_ leads to the emergence of oscillatory activity through a SNIC bifurcation (see text). Further increases in *φ*_*e*1_ result in a Fold of Limit Cycle (FLC) bifurcation, followed by a sequence of supercritical Hopf bifurcations (HB^+^). Different oscillatory regimes are marked by asterisks (*\**) and are plotted in panels (c-d), (e-f) and (g-h). Although the three oscillatory regimes are periodic, each one exhibits a different main frequency: in (c-d) *θ ≈* 4 Hz, in (e-f) *α ≈* 10 Hz, and in (g-h) *γ ≈* 40 Hz. Other parameter values are given in Table 1.

Stable (unstable) fixed points are shown in dark (light) grey, together with the amplitude of stable (unstable) periodic solutions in dark (light) colors. The local stability of periodic solutions is given by the Floquet multipliers, which determine if perturbations to such solutions grow or decay. The bifurcations related to the emergence of oscillatory dynamics are marked by vertical lines in panels (a) and (b). These figures show that the system has one stable fixed point for small inputs to *P*_1_. This stable fixed point, located on the lower branch, has three distinct pairs of complex eigenvalues and two real eigenvalues. This implies that when perturbed, the system’s response will be a combination of oscillatory behavior and exponential decay toward the steady state, in line with the results in Fig. 3.

Increasing *φ*_*e*1_ leads the system through a saddle-node on an invariant cycle (SNIC) bifurcation at *φ*_*e*1_ *≈* 107, as two fixed points collapse and a stable invariant cycle appears. Notice that this bifurcation is also encountered in the single Jansen’s model [43]. The amplitude of the oscillations decreases and the frequency increases after the system passes through two fold (saddle-node) bifurcations of limit-cycles (FLC_1,2_) at *φ*_*e*1_ *≈* 159, which are too close to each other to be seen separately in the diagram. This oscillatory activity vanishes at a supercritical Hopf bifurcation 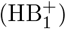 at *φ*_*e*1_ *≈* 367. Increasing *φ*_*e*1_ leads to a second supercritical Hopf bifurcation 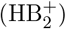 at *φ*_*e*1_ *≈* 457 and to a second invariant cycle, which has a higher frequency a much smaller amplitude (in *P*_1_) than the first one. In what follows, we analyze the oscillation dynamics of the model part by part.

We divide Figs. 5 (a) and (b) into three different regions marked with *\**’s, each one associated with different oscillatory behaviors exhibited by the system. Each one of them is shown in panels (c-g) for *v*_*P*1_ and (d-h) for *v*_*P*2_. The activity in all three regions is periodic and stable, as confirmed by the corresponding Floquet multipliers obtained with AUTO-07p [55].

In the region labeled as (*\**) in Fig. 5(a), between the SNIC and the FLC bifurcations, the system displays theta rhythmic activity with frequency *≈* 4 Hz. It is important to notice that although the main frequency of both signals is the same, *v*_*P*2_ exhibits a fast oscillation on top of the main one. In the region between FLC and the 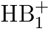, labeled as (*\*\**) in Fig. 5(a), the system exhibits alpha rhythmic activity with frequency *≈* 10 Hz, and similarly to the region before, *v*_*P*2_ shows a fast oscillation for *v*_*P*2_ superimposed on the slower one.

The results described above reveal that not only 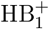 exhibits a fast oscillation, but this oscillation is locked with the phase of 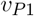. This can be understood by considering the two parts of the model, the JR and the PING, as two coupled oscillators. In regions *\** and *\*\**, the JR drives the PING activity. This driving is modulated by the 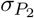 function (Eq. (3)), which increases considerably as 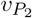 approaches *v*_0_ = 1.

Lastly, gamma rhythmic activity is observed in the region labeled as *\* * \**, after the 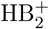 bifurcation, with frequency *≈* 40 Hz. This can be explained by analyzing the dynamics of the single Jansen’s model [43]. For this input value, the system does not oscillate; instead, it settles into a stable fixed point. In contrast, the PING model exhibits oscillations. As a result, the JR model is being driven by the PING model, but its inherent dynamics suppress the oscillations, trying to pull the system toward its fixed point and hence leading to small amplitudes in the *P*_1_ oscillations. Summarizing, for *φ*_*e*2_ = 0 the system does not exhibit simultaneously multifrequency activity, in the sense that both *v*_*P*1_ and *v*_*P*2_ share a single dominant frequency. That is, only one main frequency is observed across both populations. In the following, we analyze how increasing *φ*_*P*2_ alters this scenario and can lead to the emergence of distinct dominant frequencies for each population.

The effect of a non-zero input in the *P*_2_ population is shown in Figure 6, which presents different bifurcation diagrams with increasing *φ*_*e*2_ from left to right, using the same color scheme as Fig. 5. As we see in the first column (panels a,d), increasing the value of *φ*_*e*2_ to 20 makes little difference in the position of the first bifurcations (SNIC, FLC, and 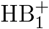). However, the bifurcation point 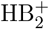 has moved significantly to the left. As we increase *φ*_*e*2_ = 40, we notice a change in the sequence of bifurcations. Now, after the FLC bifurcations, the system goes through a subcritical Hopf bifurcation 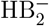 (the former 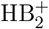 bifurcation becomes subcritical when it crosses 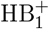), which leads to an unstable periodic solution.

**Figure 6.**
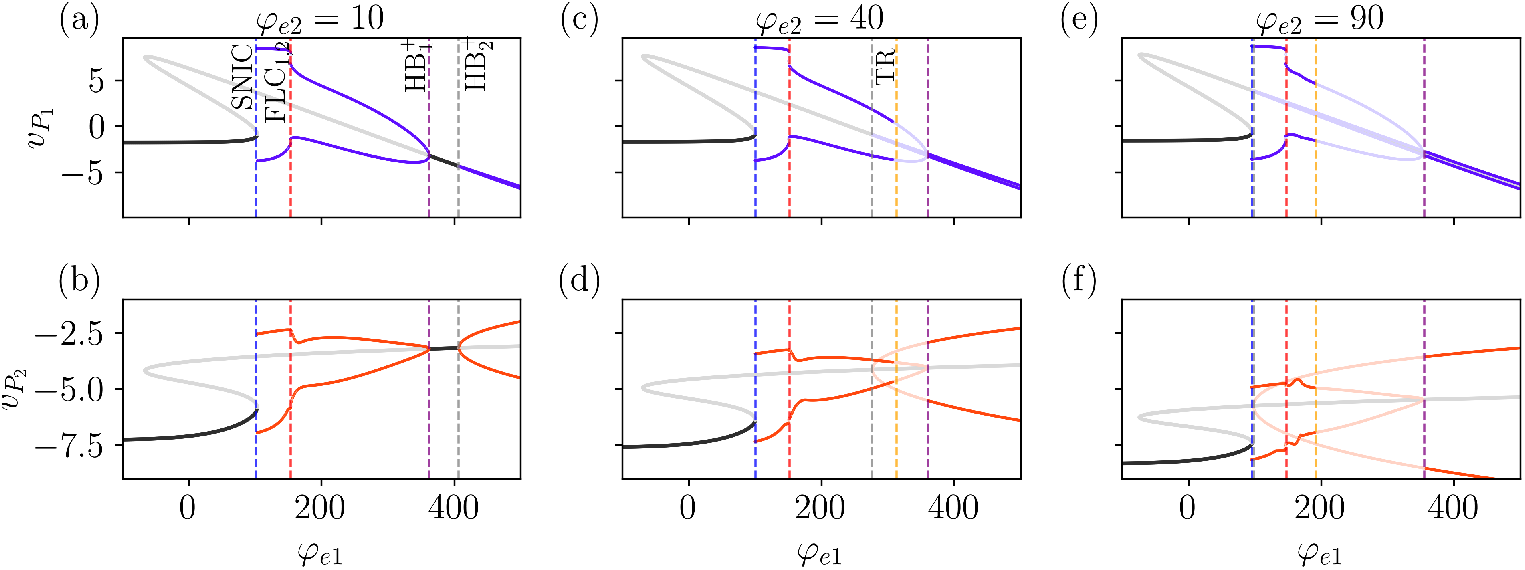
Effect of increasing *φ*_*e*2_ on the bifurcation diagram. Bifurcation diagrams obtained by varying *φ*_*e*1_ for different values of *φ*_*e*2_. Increasing *φ*_*e*2_ (left to right) lowers the threshold for the second Hopf bifurcation, 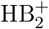, which appears at smaller values of *φe*1. Additionally, it also induces a torus bifurcation (TR). Other parameter values are given in Table 1.

Notice that from this point on, until the system reaches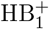, there is an overlap between former regions *\*\** and *\*\*\**, sustaining alpha and gamma rhythms, respectively. Between these two bifurcations, we notice another one, namely a torus (or secondary Hopf) bifurcation, TR (yellow dashed line in panel (c-f)). Before the TR bifurcation, the unstable inner limit cycle repels the trajectories that are attracted by the outer stable limit cycle (panel (c)). The crossing of the TR bifurcation leads to the loss of the stability of the periodic solution, leading to the emergence of the quasiperiodic behavior. In this regime, the trajectories are attracted to a two-dimensional torus and repelled from the inner limit cycle. For *φ*_*e*2_ = 90, the scenario remains qualitatively similar, with both HB^−^_2_ and TR shifting significantly to the left, which enables a larger region of overlap between the different limit cycles. Notice that the values used in Fig. 4 lie on the region slightly after the TR bifurcation.

The two-parameter bifurcation diagram (*φ*_*e*1_ *× φ*_*e*2_) shown in Fig. 7 gives us a better view of how the bifurcation points change for different parameter values. We see that except for the TR bifurcation, the other bifurcations are not significantly affected by the increase of *φ*_*e*2_, in agreement with the results of Fig. 6. To better understand how the bifurcation structure affects the frequency of the signals 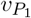 and 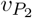, we numerically solved Eq. (5) varying *φ*_*e*1_ and *φ*_*e*2_, with results in presented in Fig. 7 panels (b-c). By comparing the main frequencies of 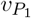 and 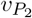 (panels (b) and (c), respectively) we verify the presence of regions where both populations display the same frequency, such as before the SNIC and after the 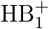 bifurcations, and also regions with multifrequency activity.

**Figure 7.**
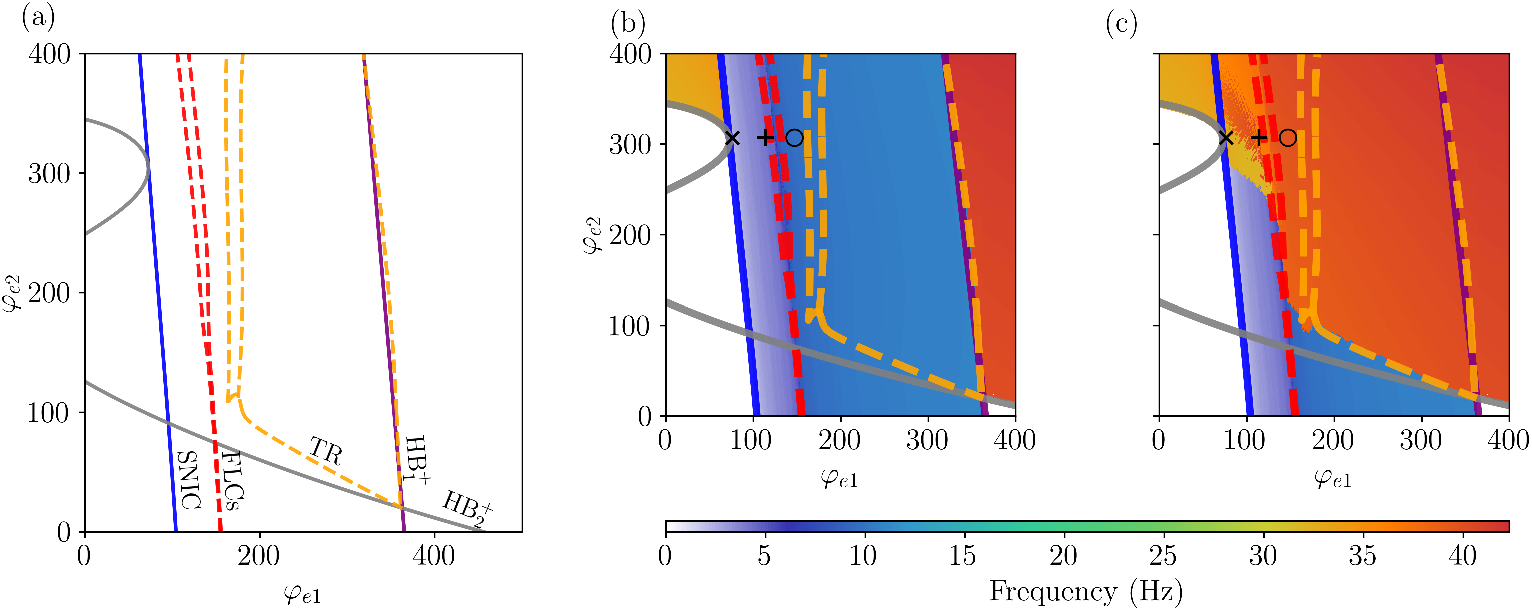
Two-parameter bifurcation diagram of Eq. (5). The color scheme used for bifurcations is the same as in Fig. 2. Panels (b) and (c) show regions colored according to the dominant frequencies of (b) 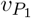 and (c) 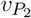, highlighting the model’s ability to sustain multifrequency activity depending on the combination of *φ*_*e*1_ and *φ*_*e*2_. The area of multifrequency activity is bounded by the SNIC (blue) and 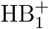 (purple) bifurcations. In panel (b), we observe that the dominant frequency of 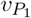 remains robust as *φ*_*e*2_ increases. Between the SNIC and FLC bifurcations, 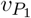 exhibits oscillations in the delta (0.5–4 Hz) and theta (4–7 Hz) ranges. As we move from the FLC to the 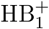 bifurcations, 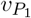 transitions to oscillations in the alpha range (8–13 Hz). For 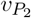 (panel c), the scenario differs. As *φ*_*e*2_ increases, the system undergoes a TR bifurcation, shifting the regime and causing the main frequency to move from the alpha range to the gamma range (*>* 30 Hz) between the FLC and 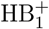 bifurcations. Additionally, in the region between the SNIC and FLC bifurcations, increasing *φ*_*e*2_ induces a change in the rhythmic activity of 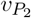, transitioning from delta/theta frequencies to gamma frequencies for *φ*_*e*2_ *>* 250. Other parameter values are given in Table 1.

Specifically, we identify two regions of multifrequency activity. The first occurs for *φ*_*e*2_ *>* 250, bounded by the SNIC and FLC bifurcations. In this region, the system exhibits frequencies ranging from approximately 2.5 to 5.0 Hz in 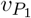, spanning both *δ* (0.5–4 Hz) and *θ* (4–8 Hz) bands, depending on the value of *φ*_*e*1_. Concurrently, 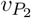 displays *γ* band activity. The second region of multifrequency activity arises as *φ*_*e*1_ increases beyond the FLC bifurcation. Here, 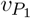 transitions into the alpha band until it reaches the 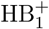 bifurcation, while 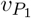 maintains *γ* frequencies (above 30 Hz) after crossing the TR bifurcation. This results in a second distinct region where the two populations oscillate at separate dominant frequencies.

These different dynamics are highlighted in Fig. 8, where the time evolution of the system corresponding to the (*φ*_*e*1_, *φ*_*e*2_) values indicated by the markers (*×*, +, ∘) in Fig. 7. Panels (a) and (b) of Fig. 8 show that *δ* activity is displayed by *P*_1_, while *γ* activity is observed in *P*_2_. Increasing *φ*_*e*1_ results in an increase of the frequency of 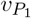 to the *θ* range, and while this component is also observed in the signal of 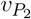, the main frequency for this populations remains on the *γ* range, as shown in panels (c) and (d). Lastly, by increasing *φ*_*e*1_ further, we show the coupling between *α* and *γ* frequencies in panels (e) and (f). Interestingly, we notice that the amplitude of 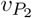 which displays *γ* rhythm is modulated by the cycle of 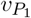 in all three cases (*δ, θ*, and *α* rhythms).

**Figure 8.**
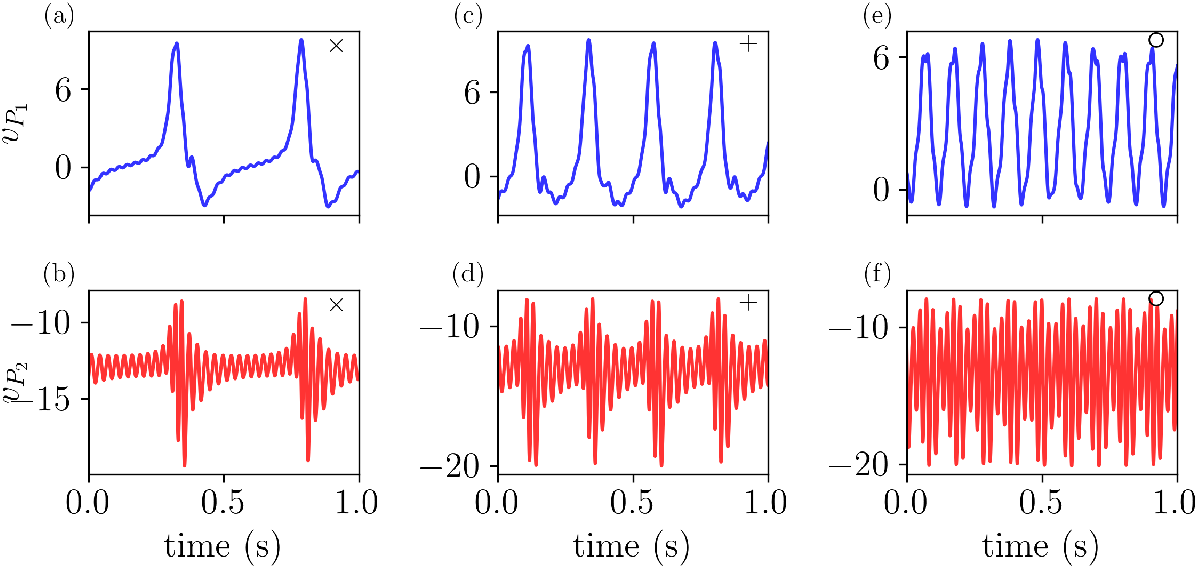
The model exhibits different frequency couplings depending on *φ*_*e*1_ and *φ*_*e*2_. The time evolution of 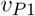 and 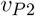 is shown for the parameter values highlighted in Fig. (7), illustrating: (a-b) coupling between *δ* and *γ* frequencies, (c-d) coupling between *θ* and *γ* frequencies, and (e-f) coupling between *α* and *γ* frequencies. All parameter values are specified in Table 1.

We further validate our results by calculating the two largest Lyapunov exponents (LEs) (see Methods), *λ*_1_ and *λ*_2_, as a function of the external inputs *φ*_*e*1_ and *φ*_*e*2_. The results are shown in Fig. 9, color-coded according to the values of (*λ*_1_, *λ*_2_), overlapped with the two-parameter bifurcation diagram. Together, these two values provide insight into the system’s dynamics: The presence of a positive LEs signifies chaos, while negative LEs indicate steadystate dynamics. If one LE is zero, the system exhibits periodic behavior. In contrast, if the system has two zero LEs, it evolves on a two-dimensional invariant torus, indicating quasiperiodicity. As expected, both periodic and quasiperiodic dynamics are observed, with the quasiperiodic regions emerging and being enclosed by torus bifurcations. Interestingly, the system also displays chaotic dynamics in the vicinity of a complex region between FLC bifurcations for high values of *φ*_*e*2_. Such a transition from quasiperiodic to chaotic dynamics is also known as *torus breakdown*. The presence of FLC bifurcations in this region implies the emergence and annihilation of an unstable limit cycle, which can create conditions favorable for chaotic dynamics.

**Figure 9.**
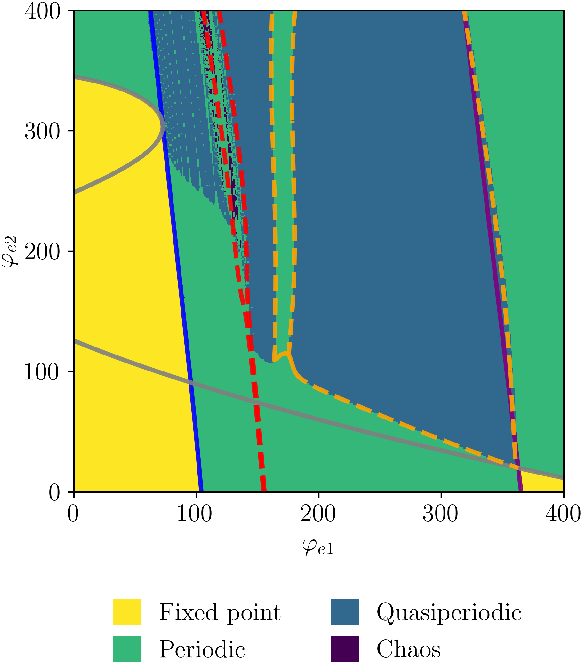
Two-parameter bifurcation diagram of Eq. (5) with regions colored according to the two largest Lyapunov exponents as a function of *φ*_*e*1_ and *φ*_*e*2_. Parameters values as in Table 1.

### 2.4 PV interneuron dysfunction

In this section we investigate the role of *PV* cells in the multifrequency observed in the previous sections, more specifically, we focus on reducing the coupling from *PV* to *P*_2_, i.e.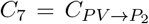. Recently, this has been proposed a mechanism to model the accumulation of amyloid-beta oligomers (A*β*O), which is supposed to damage the synaptic function of *PV* interneurons [49]. The accumulation of A*β*Os in the brain correlates with Alzheimer’s disease (AD) progression, as these soluble oligomers, formed by 2-50 monomers, are believed to be the main culprits behind various neurotoxic effects leading to cognitive decline and begin forming years before clinical signs of the disease appear [47, 56–59].

Figure 10 reproduces the two-parameter bifurcation diagram from Fig. 7, focusing on the bifurcation that defines the coexistence of limit cycles 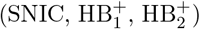 for different values of 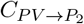. For comparison, we include the case where 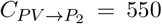, representing the baseline or healthy condition. The region where the two limit cycles coexist is highlighted in grey. Decreasing 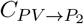 impacts all the bifurcations. For 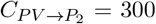 (Fig. 10(b)) and *φ*_*e*1_ = 0, all bifurcations shift slightly to the left, indicating that oscillatory activity occurs at lower levels of external inputs (hyperexcitability). The most affected bifurcation is the one related to gamma oscillations (HB^+^, grey curve), whose shift significantly enlarges the grey-shaded area, implying that multifrequency activity is easier achieved.

**Figure 10.**
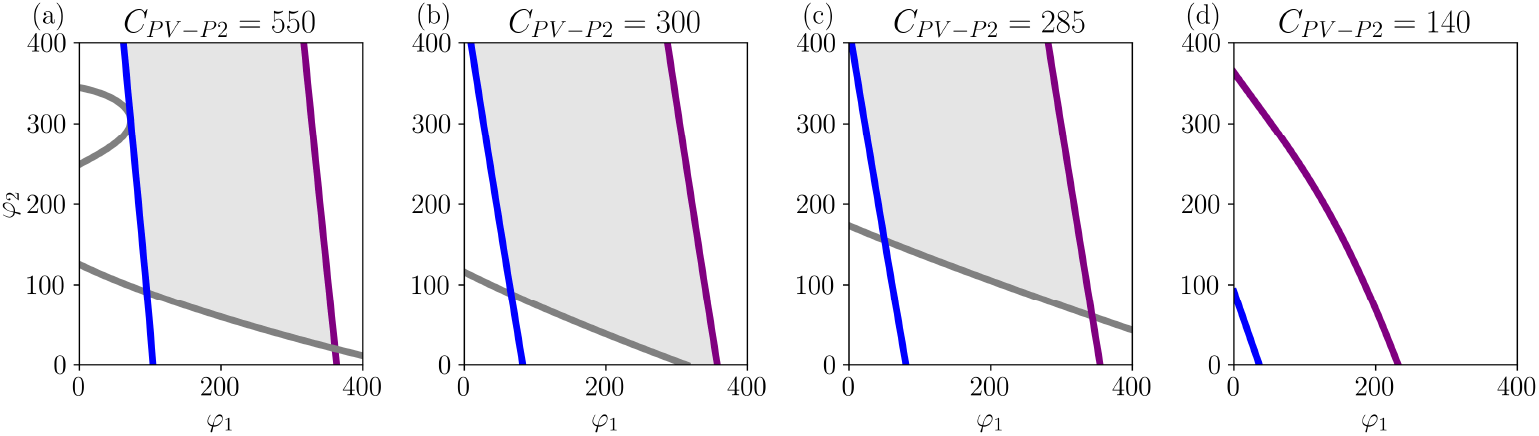
Bifurcation diagrams showing the effect of reducing 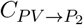 on limit cycles and gamma activity. Panel (a) shows the baseline condition, while panels (b)–(d) illustrate the progressive reduction of multifrequency activity and the eventual extinction of gamma oscillations as 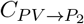 decreases. The region where the two limit cycles coexist is highlighted in grey. Parameters as in Table 1.

Further decreasing 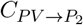 (which would corresponding to increasing the cumulative damage caused by A*β*Os) reduces gamma activity, as shown in Fig. 10(c). While the SNIC and 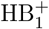 bifurcations shift further to the left, the 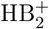 bifurcation shifts to the right, decreasing the grey-shaded area and, consequently, reducing the system’s ability to exhibit multifrequency activity.

The results described above suggest that as AD progresses, the regime of co-existence of fast and slow oscillations is reduced, with fast oscillations being disrupted the most. This is important because there is a natural variance of mean inputs to each (LaNMM) brain node as determined by dynamics and tractography. Nodes with lower mean input to P1 and P2 will be affected first.

This observation aligns with the evidence on reduced gamma power in AD [49]. Thus, the model supports the idea that early-stage AD is primarily driven by an imbalance between excitation and inhibition driven by PV interneuron dysfunction [49, 56, 58, 60]. Additionally, further reductions in 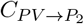 result in the complete extinction of gamma oscillations in the LaNMM model, as shown in Fig. 10(d).

### 2.5 Long-range connectivity

So far, we have considered a model with short-range connections, where all possible external inputs due to long-range connections are encoded by the parameters *φ*_*e*1_ and *φ*_*e*2_. In this section, we briefly investigate the impact of long-range connections on the dynamics of the model, particularly on its ability to sustain multifrequency activity. The coupling scheme between columns is based on the model proposed in [61] and supported by [62–64]. The final model is illustrated in the center panel of Fig. 11.

**Figure 11.**
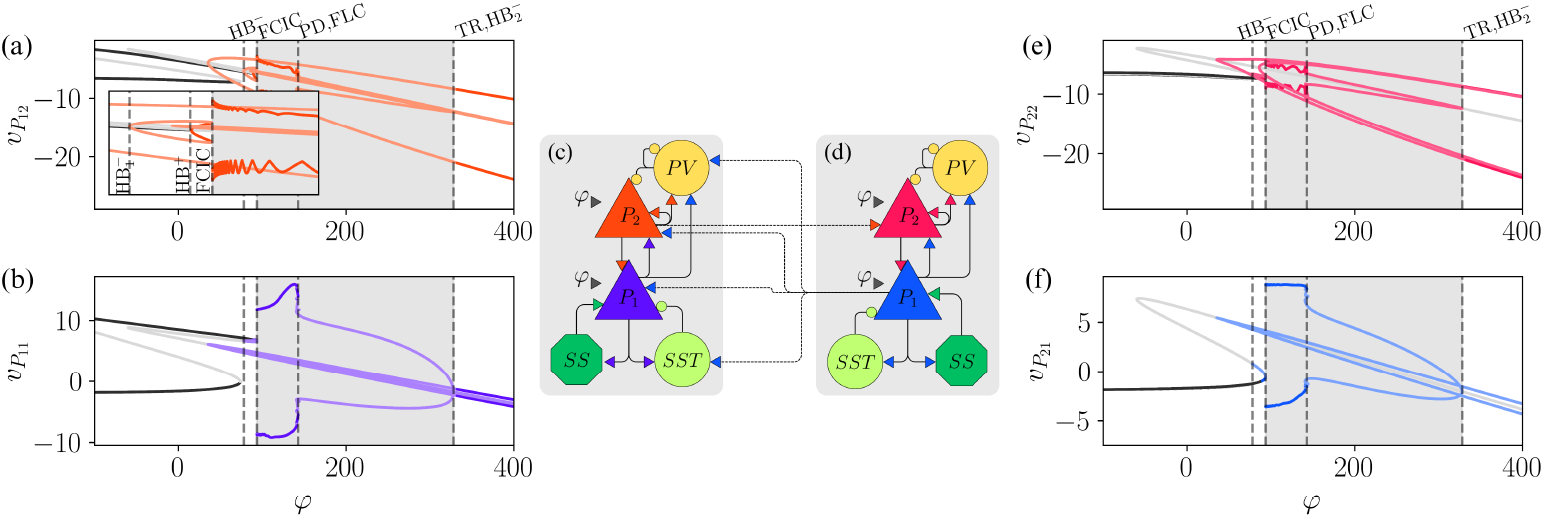
Illustration of the two-column model. (a, b) One-parameter bifurcation diagrams for the oscillators of 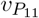 and 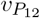 as a function of *φ*. (c, d) Schematic illustrations of the two coupled columns. (e, f) One-parameter bifurcation diagrams for 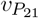 and 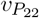 as a function of *φ*. The colored lines correspond to the respective pyramidal populations. *α* = 0.6. Two column model

This model represents two cortical columns, each consisting of two layers: a superficial layer which is modeled by the PING part of the LaNMM, and a deep layer modeled by the Jansen’s part of the LaNMM. Long-range connections represent the connectivity between the two cortical columns and are divided into feedback and feed-forward projections. Feedback projections originate from the superficial layer in column 1 and target the superficial layer in column 2. Feed-forward projections originate from the deep layer in column 2 and target both superficial and deep layers in column 1.

For simplicity, we assume that populations of the same type have identical parameters *A* and *a* across columns. We also consider that all four pyramidal populations receive the same external input, *φ*. The equations governing the final two-column model are provided in the Methods section. Building on the analysis from previous sections, we now examine how external inputs to the pyramidal populations influence the stability of fixed points and the occurrence of bifurcations focusing on the average membrane potentials of the pyramidal populations.

The results are presented in panels (a,b) of Fig. 11 for the first column (c) and in panels (e,f) for the second column (d). The bifurcation diagrams corresponding to the lower layer population (*P*_1_) are shown in the bottom panels, while those of the upper layer population (*P*_2_) are displayed in the top panels. For convenience, the populations are labeled as *P*_11_ and *P*_12_ for the first column (corresponding to *P*_1_ and *P*_2_ of the first column), and *P*_21_ and *P*_22_ for the second column (corresponding to *P*_1_ and *P*_2_ of the second column). Although the introduction of coupling between columns increases the complexity of the system compared to the single-column model, leading to richer dynamical behavior, the bifurcation scenario still bears an important resemblance with the one of Fig. 6: the coexistence of multiple limit cycles.

Before the emergence of any limit cycle in the bifurcation diagrams of Fig. 11, we notice that the system exhibits one additional stable fixed point compared to the original model, leading to bistability. The emergence of the first limit cycle happens at *φ ≈* 35 through an 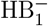 bifurcation at *φ ≈* 35, which is unstable until it goes through a TR bifurcation around *φ ≈* 332. Further increasing *φ* leads to another HB^−^ near *φ ≈* 86, where an unstable limit cycle emerges. This is followed by an HB^+^, associated with a stable limit cycle, as shown in the inset of Fig. 11(a). These two limit cycles collide in a fold of cycles bifurcation (FCIC), giving rise to a stable limit cycle. Once again, the system’s main frequency remains below *α*, while the *P*_2_ populations exhibit a *γ* component, which allows *δ*-*γ* and *θ*-*γ* couplings. As *φ* continues to increase, the system undergoes a series of period-doubling (PD) and FLC bifurcations around *φ ≈* 142. Following these transitions, both columns exhibit *α*-*γ* coupling. The multifrequency activity ceases when the limit cycle originating from the FCIC bifurcation disappears in an 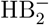 near the TR bifurcation. Beyond this point, both columns oscillate solely at the *γ* frequency.

The asymmetry in the coupling causes the dynamics of the populations in different columns to differ. For instance, compared to the original model, *P*_11_ receives an extra input from *P*_21_, and two populations projecting into *P*_11_ (*P*_12_ and *SST*_11_) also receives an input from *P*_21_. Across the range of *φ* studied, *P*_11_ exhibits a larger amplitude than *P*_21_, which reflects the larger number of pyramidal neurons being activated in *P*_11_. For the top layer pyramidal populations, *P*_12_ and *P*_22_, although the amplitudes are similar, the dynamics for the range between the SNIC and the FLC bifurcations is rather different.

The time evolution of the membrane potentials is shown in Fig. 12. The top row displays the dynamics of 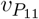 and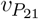, while the bottom row shows 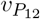 and 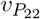. As the external input *φ* increases from *φ* = 100 (a–b) to *φ* = 125 (c–d) and *φ* = 190 (e–f), the activity of the *P*_1_ populations shifts from the delta to the alpha frequency band. A similar trend is observed in 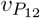 and 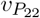, although with a more pronounced gamma component. This indicates that while the coupling between columns adds complexity to the model, as observed in the bifurcation diagram, the core feature of the model remains robust and is governed by the same mechanism: the coexistence of limit cycles.

**Figure 12.**
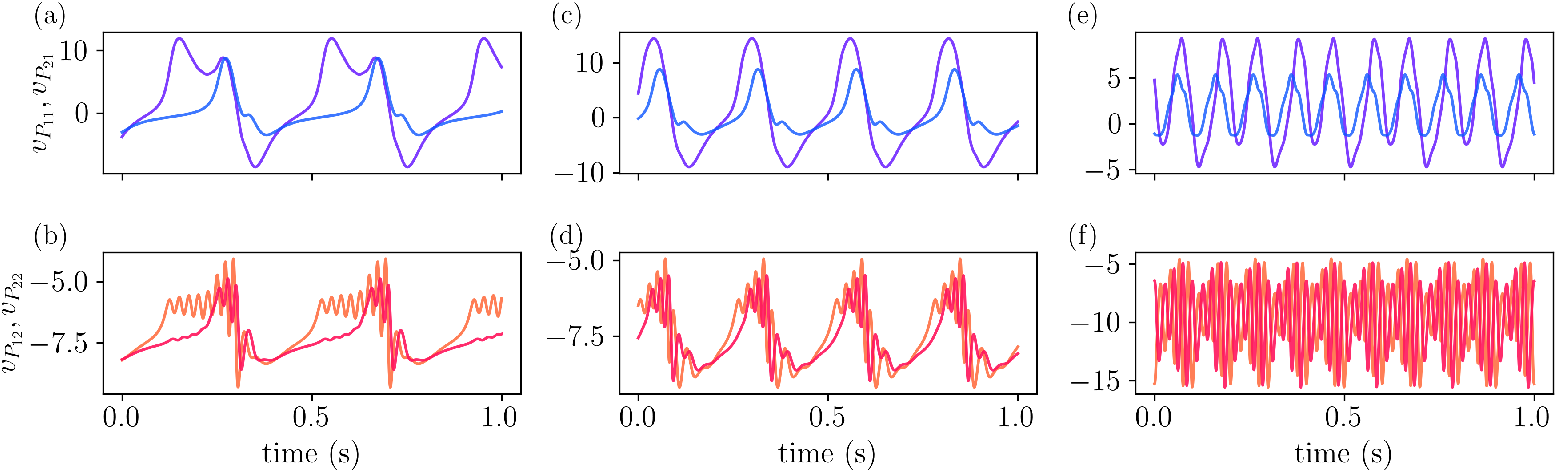
Time evolution of membrane potentials in the two-column model. Top row: Dynamics of 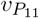 and 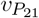 as the external input *φ* increases, illustrating a transition from delta to alpha frequency bands. Bottom row: Corresponding dynamics of 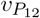 and 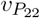, showing similar trends but with a prominent gamma-band component. (a–b): *φ* = 100; (c–d): *φ* = 125; (e–f): *φ* = 190.

## 3 Discussion

In this paper we investigate the dynamical properties of a laminar neural mass model, designed to capture different dynamics across cortical layers. The model achieves this by combining two neural mass models, each exhibiting oscillatory dynamics in distinct natural frequency bands, meaning that under external perturbations, each component oscillates at different frequencies. As in the original work, we consider that both pyramidal populations are targeted by external inputs *φ*_*e*1_ and *φ*_*e*2_. To analyze the system’s behavior, we first use *φ*_*e*1_ as the bifurcation parameter while keeping *φ*_*e*2_ = 0, allowing us to examine its role in triggering oscillatory dynamics. As shown in Fig. 5, the oscillatory regime can be divided into three regions depending on the value of *φ*_*e*1_, with the dominant frequency transitioning from *θ* to *γ*. Additionally, for intermediate values of *φ*_*e*1_, the oscillatory behavior resembles that observed in the Jansen model, whereas for higher values, the activity accelerates, corresponding to a PING mechanism for gamma activity.

We also investigate the role of *φ*_*e*2_ in the emergence of rhythmic activity. This input directly targets the PING component of the model, and thus we expect it to facilitate the emergence of gamma oscillations. This is precisely what we observe in Fig. 6: as we increase the value of *φ*_*e*2_, the HB bifurcation that gives rise to gamma activity occurs at smaller values of *φ*_*e*1_. This clarifies the mechanism behind the multifrequency activity observed in LaNMM: the coexistence of limit cycles, one sustaining low-frequency oscillations and the other supporting high-frequency oscillations. We verify the robustness of this mechanism through a two-parameter bifurcation analysis as a function of *φ*_*e*1_ and *φ*_*e*2_, shown in Fig. 7, where we observe a large region in the parameter space where the two limit cycles coexist.

Of particular interest are the regions delimited by the torus bifurcations (TR), where the system exhibits quasiperiodic dynamics, and both *α* and *γ* frequencies have significant power. Within this region, the model is capable of exhibiting multifrequency activity across different frequency ranges, including *δ*-*γ, θ*-*γ*, and *α*-*γ*. We conclude our analysis by evaluating the Lyapunov exponents of the model, shown in Fig. 9. While quasiperiodicity is observed over a large area of the parameter space, we also identify periodic dynamics in regions where both limit cycles coexist. This behavior can be understood by considering the two layers of the model as entrained oscillators. In such scenarios, the oscillators can resonate, a phenomenon that gives rise to Arnold tongues. This is precisely what we observe in the regions of periodic activity. Moreover, we also observe chaos small regions in the parameter space, in line in recent results.

We highlight the significance of the region where the system exhibits two limit cycles by modeling PV interneuron dysfunction, recently proposed as a mechanism to represent A*β*O, a key biomarker of Alzheimer’s disease progression. As PV cell connectivity deteriorates, the system gradually loses its ability to generate multifrequency activity and, in advanced cases, ceases to oscillate altogether. This suggests that as the disease progresses, the coexistence of fast and slow oscillations diminishes, with fast oscillations being the most affected, in line with the results in [49].

We also extended the model to account for long-range connectivity. By incorporating a feedforward and feedback coupling scheme between columns, we demonstrate that the coexistence of limit cycles persists under these conditions. This is confirmed through a bifurcation analysis of the extended model, as shown in Fig. 11. As expected, this model exhibits a more complex bifurcation structure; however, more importantly, rhythmic activity emerges in both the bottom and top layers. As a result, multifrequency activity is observed in both columns at different frequency ranges, as illustrated in Fig. 12.

We emphasize that the mechanism of coexisting limit cycles underlies the approach of incorporating different neural populations to account for multiple frequency ranges in neural mass models, a concept that is not new. The strength of the LaNMM model lies in its ability to exhibit diverse frequency couplings using only five neural populations, controlled solely by adjusting the level of external inputs. While previous models have also been capable of generating multiple frequency interactions, they typically required either increasing the number of neural populations or modifying the time constants of specific populations (*a* and *φ*_0_ in our case) to access different frequency regimes [41, 65, 66]. This demonstrates the efficiency of LaNMM in capturing complex neural dynamics with a relatively simple yet powerful parametrization. In order to further refine the LaNMM, future work should generalize our formalism towards next-generation neural mass models [9]. These modeling framework provides additional equations, rigorously derived from microscopic dynamics, for each neural population mean firing-rate and membrane potential, to which the sigmoid function employed here represents a quasi-stationary approximation [67, 68]. To the best of our knowledge, multifrequency activity in these models has been studied only for the case of external forcing [69], and two interacting inhibitory populations with different time scales [70].

Our analysis highlights the potential of the LaNMM to model various phenomena and different brain regions in future studies. The ability to exhibit multifrequency activity is a prerequisite for a model to capture cross-frequency coupling phenomena such as phase-amplitude coupling (PAC) or amplitude-amplitude coupling (AAC) and, features commonly observed in neural recordings. For instance, the three frequency couplings observed here are also found in brain activity: *δ*-*γ* coupling has been proposed as a biomarker of postictal activity [71] and is linked to dopamine modulation [72]; *θ*-*γ* coupling is observed in the hippocampus of both rodents and humans [73, 74]; and *α*-*γ* coupling has been associated with cognitive processes [36, 75, 76]. Recent results show that the cross-frequency coupling exhibited by the LaNMM can provide the key mechanisms for predictive coding, gating [33] and cooperation/competition across brain regions [34].

## 4 Methods

### 4.1 Single column model

Following the diagram shown in Fig. 1 (a) of [22], the LaNMM is given in the synapse-driven formalism by

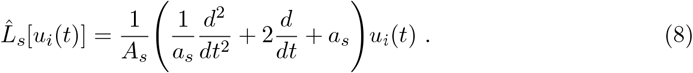

With

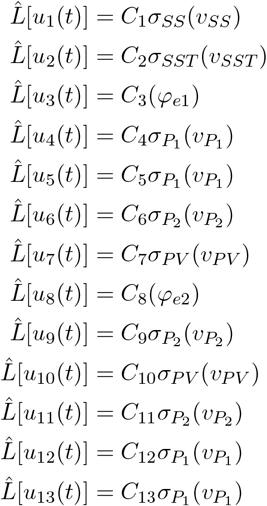

Where *s* is the type of neurotransmitter and *i* the population in question. Notice that 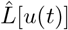 has a dimension of frequency (Hz). The average membrane potentials are given by:

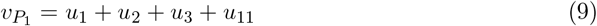

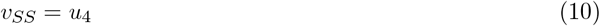

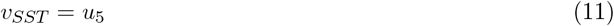

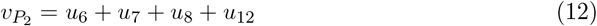

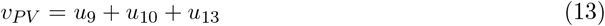

Using the the above equations and taking out the repeated synapses, and since all the synapses coming from one population have the same dynamics and the only difference is the coupling strength relative to the targeting population, we get

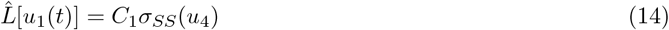

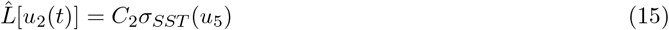

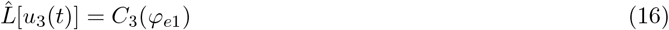

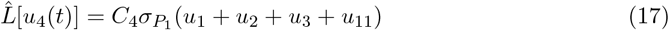

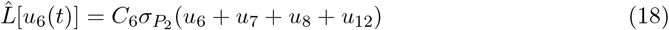

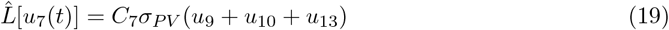

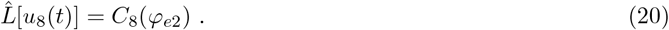

Notice that *u*_4_, *u*_5_, *u*_12_, and *u*_13_ all represent the perturbation caused by *P*_1_, which will be weighted by the average number of connections between *P*_1_ and the targets, *u*_4_. Similarly, *u*_6_ = *u*_9_ = *u*_11_ and *u*_7_ = *u*_10_. Using these identities we can write:

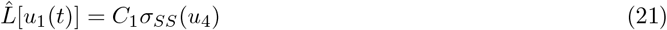

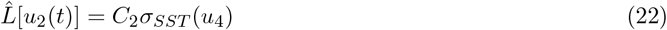

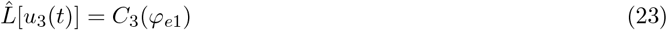

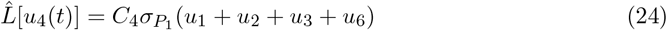

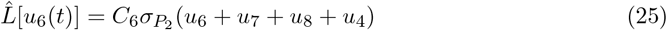

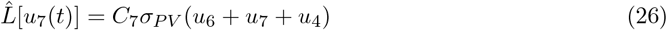

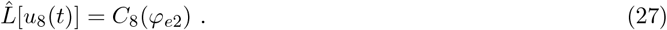

For the sake of coherence with the main text we will introduce the change of variables:

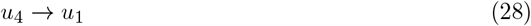

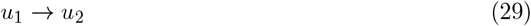

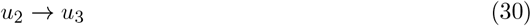

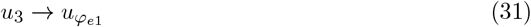

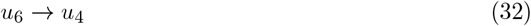

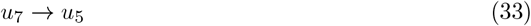

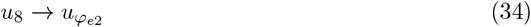

which gives us:

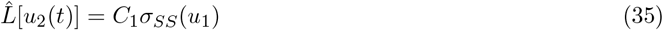

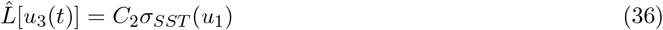

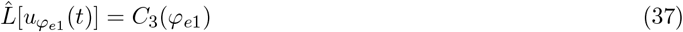

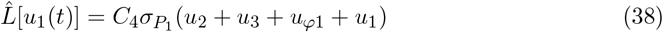

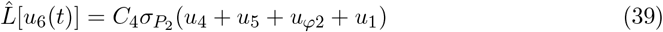

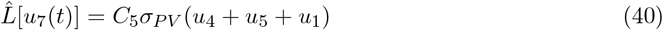

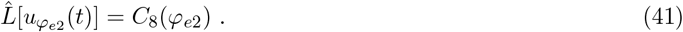

We now introduce a change of variables *u*_*i*_ = *C*_*j*_*y*_*i*_, where *C*_*j*_ accounts for the connectivity of the incoming synapse. The first equation, represents the process of receiving an average synapse and producing a membrane perturbation from the *point of view* of the *SST* population. The incoming average synapse from *P*_1_ is weighted by *C*_4_ and the outgoing average synapse to *P*_1_ is weighted by *C*_1_. Taking this into account we have

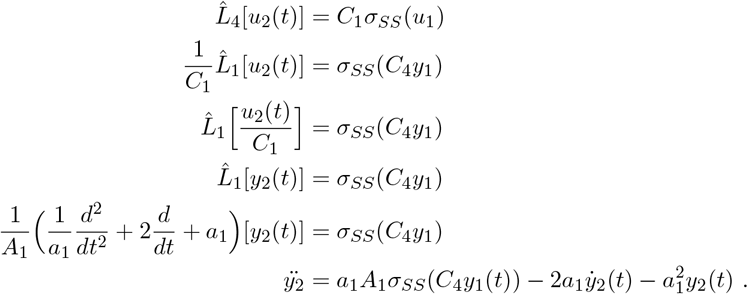

For the external inputs, we have

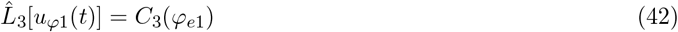

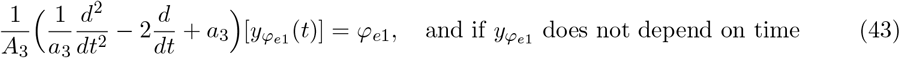

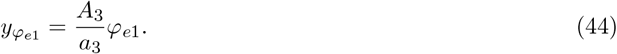

Similarly, for *s*_8_ we have 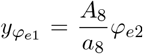.Putting these results together and using the correspondent parameters for *A*’s and *a*’s we have:

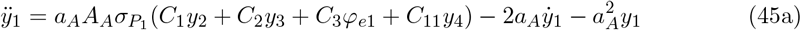

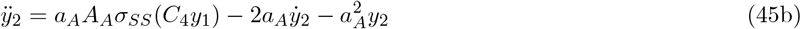

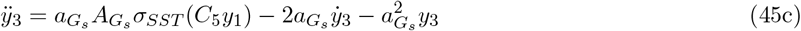

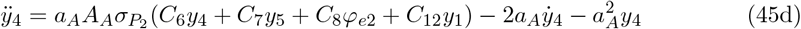

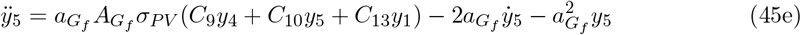

where each one of these second-order equations can be rewritten as a system of two first-order equations using 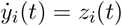.

### 4.2 Two-column model

Following the diagram shown in Fig. 11 (c,d) the equations of the two column model are given by:

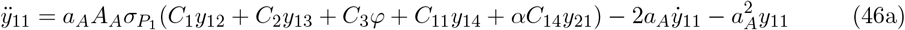

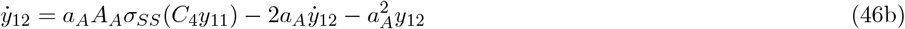

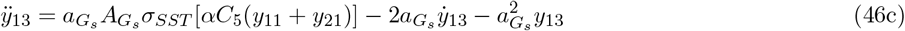

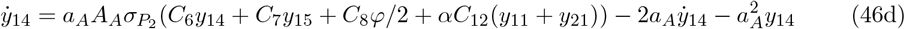

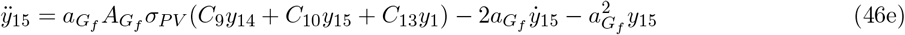

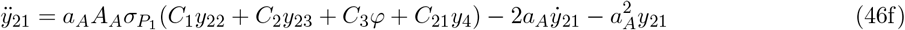

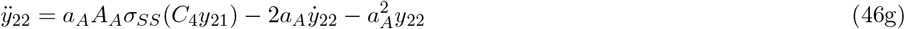

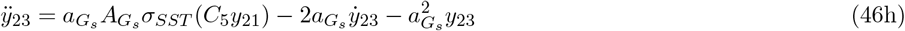

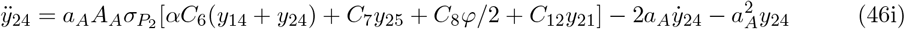

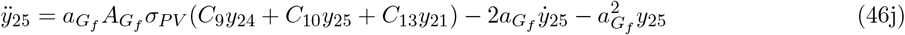

To balance the influence of additional inputs on the populations, we introduce the parameter *α*, which controls the strength of these inputs. For instance, in Eq. (46c), the *SST*_11_ population receives excitatory inputs from both *P*_11_ and *P*_21_, and *α* modulates the contribution of these inputs. Additionally, we introduce *C*_14_, which represents the number of synaptic connections between the *P*_21_ and *P*_11_ populations. All numerical simulations and bifurcation diagrams were performed with *α* = 0.6 and *C*_14_ = 56.25. Note that *α* = 0.0 corresponds to the case where the two columns are uncoupled. The impact of this parameter on the system’s dynamics will be explored in future work.

### 4.3 Numerical simulation and frequency analysis

Equations (5) were numerically solved using Python’s scipy.solve ivp function [77], employing a fourth-order Runge-Kutta algorithm, over a 100-second duration after discarding a 10-second transient, with an integration step of *dt* = 0.001. The frequency analysis for Fig. 4 was performed using Python’s `scipy.fft.fft` [77], with 1-second windows, proper windowing and padding, and averaging. For computational efficiency, in Fig. 7, the entire signal was used with windowing and padding, and the peak frequency was obtained for *P*_1_ and *P*_2_.

### 4.4 Bifurcation diagrams

All bifurcation diagrams were calculated with AUTO-07p, a publicly available software for continuation and bifurcation problems [55].

### 4.5 Lyapunov exponents

Lyapunov exponents are commonly used to assess how perturbations affect the trajectory of a system over time. A system has as many Lyapunov exponents as its dimensions, but by focusing on the two largest, *λ*_1_ and *λ*_2_, we can classify the system’s dynamics as follows:

1. Fixed points: both exponents negative (*λ*_1_, *λ*_2_ *<* 0).
2. Periodic dynamics: one zero exponent, the rest negative (*λ*_1_ = 0, *λ*_2_ *<* 0).
3. Quasiperiodic dynamics: both exponents zero (*λ*_1_ = 0, *λ*_2_ = 0).
4. Chaotic dynamics: one positive exponent (*λ*_1_ *>* 0, *λ*_2_ *≤* 0).

The Lyapunov exponents for Eq. (5) were calculated using the lyapunovspectrum function from the ChaosTools module in Julia, part of the DynamicalSystems library [78, 79]. A threshold of |*λ*_*k*_| *<* 10^−3^ was applied to differentiate between zero and non-zero values.

